# High selfing capability and low pollinator visitation in the epiphyte *Pitcairnia heterophylla* (Bromeliaceae) at a Costa Rican cloud forest

**DOI:** 10.1101/057109

**Authors:** Luis D. Ríos, Alfredo Cascante-Marín

## Abstract

Most epiphytic bromeliads exhibit specialized pollination systems likely to promote out-crossing but, at the same time, possess floral traits that promote autonomous selfing. Adaptations that promote selfing in flowering plants with specialized pollination systems have been considered as a mechanism for reproductive assurance. In this paper, we analyzed the breeding system and pollinator visitation rate of the hummingbird-pollinated bromeliad *Pitcairnia heterophylla* in order to see if they fit such trend. We performed hand pollination experiments, video recording of floral visitors, and recorded floral traits in order to describe the reproductive and pollination system of the studied species in a cloud forest in Costa Rica. Results from the pollination treatments indicated that *P. heterophylla* is self-compatible (SCI_f_ = 0.77), capable of autonomous pollination (AFI_f_ = 0.78), and non-agamospermous (AG_f_ = 0.01). Floral traits, such as scentless red flowers, with tubular corolla and nectar production, suggested ornithophily which was confirmed by the video recording of *Lampornis calolaemus* (Trochilidae) visiting flowers. However, the visitation rate was low (0.6 visits day-1 per plant) based on 918 hours of video recording using trail cameras. We suggest that the high selfing capability of the studied population of *P. heterophylla* might be related to the low pollinator visitation rate. If low pollinator visitation is common among hummingbird-pollinated and epiphytic bromeliads, then selfing could be a widespread mechanism to enhance their reproductive success.

## Introduction

Epiphytes, plants that grow on other plants without extracting nutrients nor water from them (Benzing 1990), are a taxonomically diverse and abundant group in moist tropical forests and may represent up to one-third of the local flora in some Neotropical areas (Gentry & Dodson 1987). Epiphytes represent nearly 9% of all vascular plants (ca. 27 614 species) with orchids, bromeliads, and aroids being the most important groups of flowering epiphytes (Zotz 2013).

Most angiosperm epiphytes display specialized pollination systems which involve a series of particular floral traits and pollen vectors such as hummingbirds, bats, and bees, in a relationship intended to promote pollen flow among conspecifics and an outcrossing mating system (Ackerman 1986; Benzing 1987). However, a general trend among flowering plants with specialized pollination systems is the parallel presence of reproductive traits that also facilitate self-pollination (Fenster & Marten-Rodriguez 2007); which suggest the chance for selfed progeny and a mixed mating system (Goodwillie et al. 2005). Specialized pollination systems in epiphytes may enhance pollen flow between conspecific plants but the additional presence of mechanisms that promote autogamy may compensate for the reduced capacity of epiphytes to attract pollinators as a consequence of their highly aggregated spatial distribution in the forest and low floral displays (Bush & Beach 1995). Levin (1972) hypothesized that, in circumstances of competition for pollinator service, a selective response to reduce the reliance upon such service might favor self-pollination mechanisms to accomplish reproductive assurance.

Epiphytic bromeliads (Bromeliaceae) are highly diverse and almost exclusive to Neotropical areas (Benzing 2000), where nearly 3160 epiphyte bromeliads (56% of the family) inhabit the forest canopy (Zotz 2013). Epiphytic bromeliads are frequently pollinated by hummingbirds and bats (Benzing 2000). Thus far, the breeding system of only 2.5% of the known epiphytic bromeliads has been studied and there seems to be a high incidence of self-compatible species capable of autonomous self-pollination (reviewed by Matallana *et al.* 2010). Matallana et al. (2010) suggested that self-pollination and self-compatibility in bromeliads has evolved as a mechanism to avoid interspecific pollen flow among congeners in highly diverse ecosystems. Molecular studies regarding the mating system of epiphytic bromeliads pollinated by hummingbirds reveal a highly autogamous mating system congruent with a high ability for self-pollination (Cascante-Marín *et al.* 2006).

Self-pollination in epiphytic bromeliads might counteract pollinator unpredictability in forest canopies and enhance a higher reproductive success when pollinator visits are scarce. We evaluated this trend in *Pitcairnia heterophylla* (Bromeliaceae: Pitcairnioideae), an epiphytic (rarely saxicolous) and C3-type bromeliad (Reinert *et al.* 2003). Self-compatibility and self-pollination have been documented in the genus *Pitcairnia* (Wendt *et al.* 2001, 2002; Fumero-Caban & Melendez-Ackerman 2007; Bush & Guilbeau 2009). *Pitcairnia heterophylla* possesses floral traits that suggest a specialized pollination system involving hummingbirds, which include tubular and scentless red flowers, and nectar production (Smith & Downs 1974).

In this paper, we document the pollination and mating system of *P. heterophylla* by examining the reproductive biology of an epiphytic population in a Costa Rican cloud forest. Our research had the following objectives: 1) to describe the floral biology and nectar production, 2) to determine the plant’s breeding system using pollination treatments, and 3) to estimate pollinator visitation rates to individuals of the studied species.

## Materials and methods

### Study species and site

*Pitcairnia heterophylla* (Lindl.) Beer has a wide geographic distribution from Mexico to Venezuela and Peru, between 100-2500 m asl (Smith & Downs 1974). Ramets possess spiny-serrate modified leaves and long linear leaves that are shed during floral anthesis (Smith & Downs 1974). Individuals (i.e. genets) of *P. heterophylla* at the studied population were composed of three to 73 ramets (mean=31.4 ± 21.7 SD, N=20). Each ramet potentially develops a single and short (ca. 2.5 cm) terminal inflorescence with 5-17 hermaphroditic flowers (mean length=4.0±0.2 cm, N=20) (Fig. 1). Fruits are dehiscent dry capsules which release wind-dispersed seeds (Smith & Downs 1974). The studied population exhibits an annual flowering frequency from January to March (dry season) and seed dispersal takes place the following dry season (A. Cascante-Marín, unpubl. data). A voucher specimen was deposited at the herbarium of the University of Costa Rica *(Ríos 28, USJ)*.

**Fig. 1.**
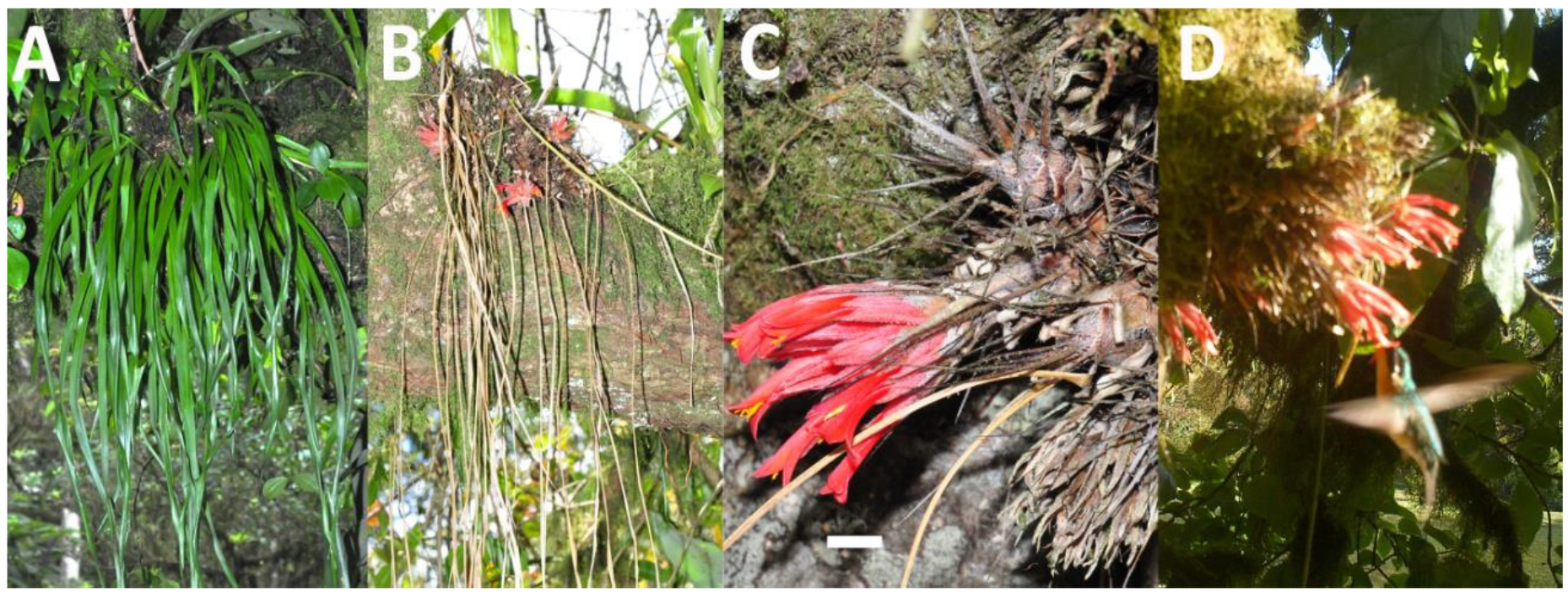
Habit of an epiphytic genet of *P*. heterophylla (Bromeliaceae) showing the long linear leaves (A). A leafless genet during the flowering period, note the short red inflorescences and the presence of dried leaves still attached to the ramets (B). Close-up of an inflorescence, note the short peduncle and the ramets formed by modified spinose leaves; scale bar = 1 cm (C). A female hummingbird, Lampornis calolaemus (Trochilidae), visiting the flowers of a P. heterophylla genet (D).

Field work was conducted in a montane cloud forest in the vicinity of the Central Valley in Costa Rica, Cartago province (reference coordinates: 9°53’20” N; 83°58 10” W), known as La Carpintera. The site consists of an elevated mountainous terrain rising from 1500 to 1800 m asl, the forested area consists of an irregular fragment of nearly 2400 ha that covers the ridge and mountain slopes and is mostly composed of old secondary forest (>50 y) interspersed with older remnant forest patches. Some representative forest trees are oaks *(Quercus* spp.), fig trees *(Ficus* spp.), and members of the avocado family (Lauraceae) among others (Sánchez et al. 2008). Mean annual precipitation is 1839.2 mm and mean annual temperature is 16.1°C. The annual distribution of rainfall follows a seasonal pattern, with a period of low precipitation or dry season (<60 mm per month) from December to April (IMN, undated).

### Breeding system

During the 2013 flowering season, controlled pollination experiments were performed on 89 flowers from 23 individuals (3-6 flowers per individual). Flowers were assigned to the following treatments: (1) Agamospermy (flowers had their stigma removed at the beginning of anthesis); (2) Autonomous self-pollination (un-manipulated bagged flowers); (3) Hand self-pollination (flowers hand-pollinated with their own pollen); (4) Hand crosspollination (previously emasculated flowers hand-pollinated with pollen from another plant), and (5) Natural pollination (un-manipulated and randomly selected flowers exposed to pollinators in the field). Pollen was removed from the anthers and applied to the stigma’s surface using a thin paint-brush. Manipulations were carried out with plants kept at a greenhouse in the study site. Fruit and seed set were evaluated ten months after the application of the pollination treatments.

The breeding system was determined by using Lloyd and Schoen’s (1992) selfcompatibility index (SCIf) and autofertility index (AFIf), and Ramirez and Brito’s (1990) index of agamospermy (AGf). SCIf was calculated by dividing the mean number of seeds per fruit in the self-pollination treatment by the number of seeds in the artificial crosspollination treatment. AFIf was estimated by dividing the mean seed number per fruit in the autonomous self-pollination treatment by the seeds produced by the hand-crossed pollination experiment. Finally, AGf was calculated as the ratio of seeds produced in the agamospermy treatment to the seeds developed in naturally pollinated flowers. Differences in seed production among treatments were statistically compared by means of a non-parametric test (Kruskal-Wallis) using the statistical software PAST v. 2.16 (Hammer *et al.* 2001), further pair-wise comparisons between treatments were performed with Mann-Whitney Tests after a Bonferroni correction (*K*=5, *P*=0.01).

### Flower development and nectar production

Inflorescence and flower development were recorded on 20 randomly chosen individuals in the field during the 2013 flowering season. For each individual, we recorded inflorescence bud emergence, and floral anthesis. Development of individual flowers was followed on one flower per inflorescence. Nectar volume and sugar concentration were analyzed on 18 unvisited flowers of six individuals at 3-h intervals from 0600 to 1800 h. Flowers were protected with a mesh bag and nectar was harvested by inserting a calibrated micropipette. Nectar concentration (% sucrose) was measured with a hand-held refractometer (Bellingham & Standley Ltd., United Kingdom) at 22-24°C.

### Flower visitors

The identity and visitation frequency of potential pollinators under natural conditions were documented with the use of trail cameras (Trophy Cam, Bushnell Corp., Overland Park, Kansas, USA). We filmed inflorescences from eight genets from January 29 to February 27 of 2013; cameras were positioned at a suitable distance (about 1 m away) to facilitate visitor’s identification. We recorded visits from 0600 to 1800 h and for an average filming time of 115 (±52) hours per plant and totaling 918 recording hours. A legitimate visit was recorded whenever a visitor contacted both pistil and stamens. The visitation rate was calculated as the mean number of visits per hour across the monitored genets.

## Results

### Breeding system

Fruit set was lower (89%) under autonomous pollination conditions compared to artificially (selfed and crossed) and naturally pollinated flowers (Table 1). Fruit set by agamospermy was minor (6.7%). Seed set significantly differed among pollination treatments (*H*=46.03, *P*<0.001). Manually cross-pollinated flowers produced the highest number of seeds (670 seeds fruit^−1^), followed by flowers exposed to natural pollination and manually self-pollinated flowers (540 and 516 seeds fruit^−1^, respectively; Table 1). The relatively high SCIf and AFIf index value (0.77 and 0.78, respectively) indicates that *P. heterophylla* is self-compatible and capable of autonomous pollination (Table 1). Agamospermy was negligible, AG_f_=0.01.

**Table I.**
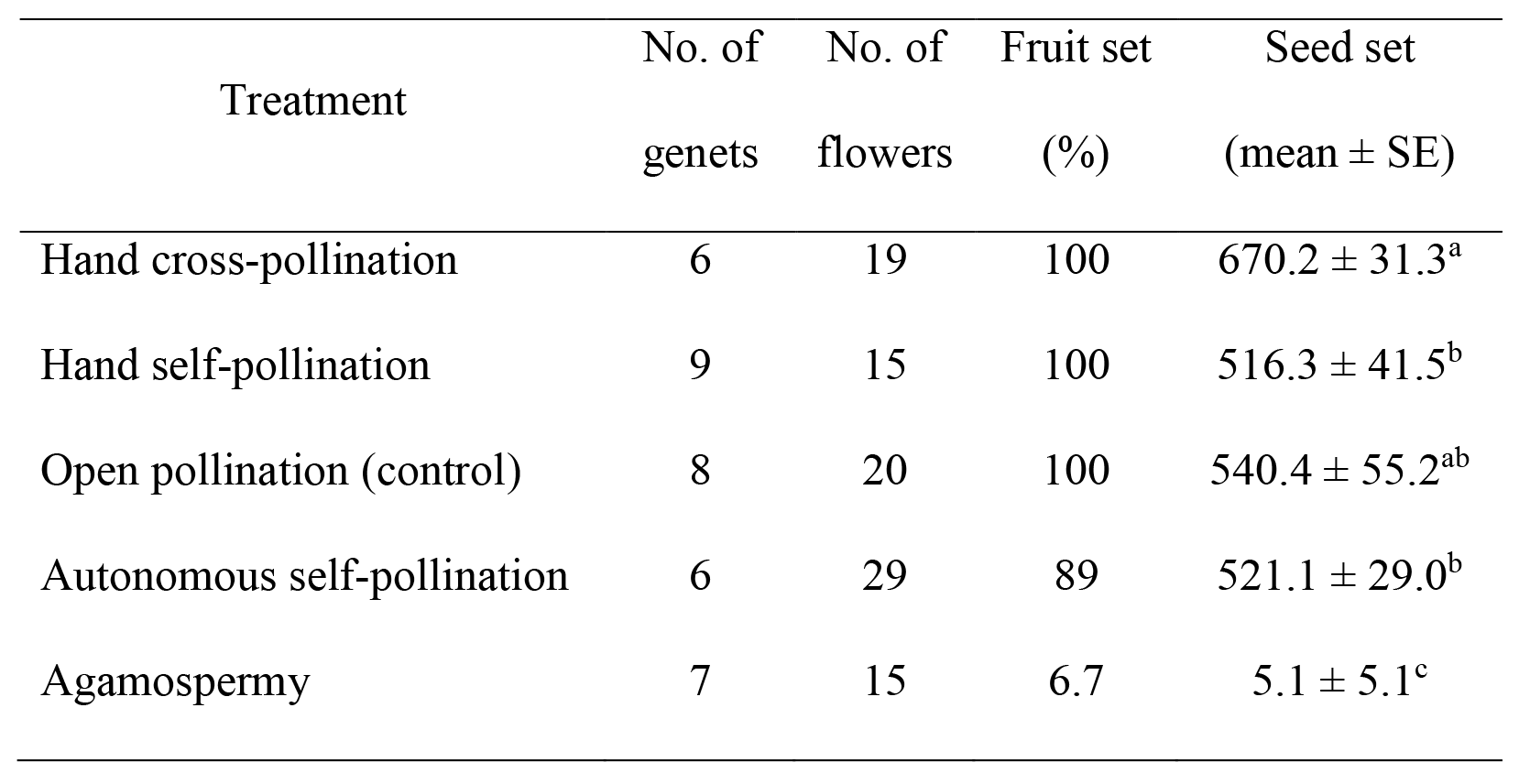
Results from natural and hand-pollination treatments on Pitcairnia heterophylla (Bromeliaceae) at a cloud montane forest, Costa Rica. Letters represent significant differences between treatments after a Mann—Whitney test (Bonferroni correction: *K*=5 *P*=0.01). N=89 flowers and 23 genets.

| Treatment | No. of<br>genets | No. of<br>flowers | Fruit set<br>(%) | Seed set<br>(mean ± SE) |
| --- | --- | --- | --- | --- |
| Hand cross-pollination | 6 | 19 | 100 | 670.2 ± 31.3<sup>a</sup> |
| Hand self-pollination | 9 | 15 | 100 | 516.3 ± 41.5<sup>b</sup> |
| Open pollination (control) | 8 | 20 | 100 | 540.4 ± 55.2<sup>ab</sup> |
| Autonomous self-pollination | 6 | 29 | 89 | 521.1 ± 29.0<sup>b</sup> |
| Agamospermy | 7 | 15 | 6.7 | 5.1 ± 5.1<sup>c</sup> |

### Floral biology and nectar production

Signs of inflorescence bud development were perceptible by early November. Inflorescences completed their development within three to four weeks and by mid-January the population flowering season had started. A single inflorescence opened flowers from one to three weeks. No signs of dichogamy or herkogamy were perceptible. Mean accumulated volume of nectar in unvisited flowers was 6.9 μL ± 0.9 (±SE) (Fig. 2A), with a mean sucrose concentration of 16.6 ± 0.6 % (Fig. 2B). Major nectar secretion and sucrose concentration occurred during early anthesis in the morning at around 0600 h (Fig. 2A, B).

**Fig. 2.**
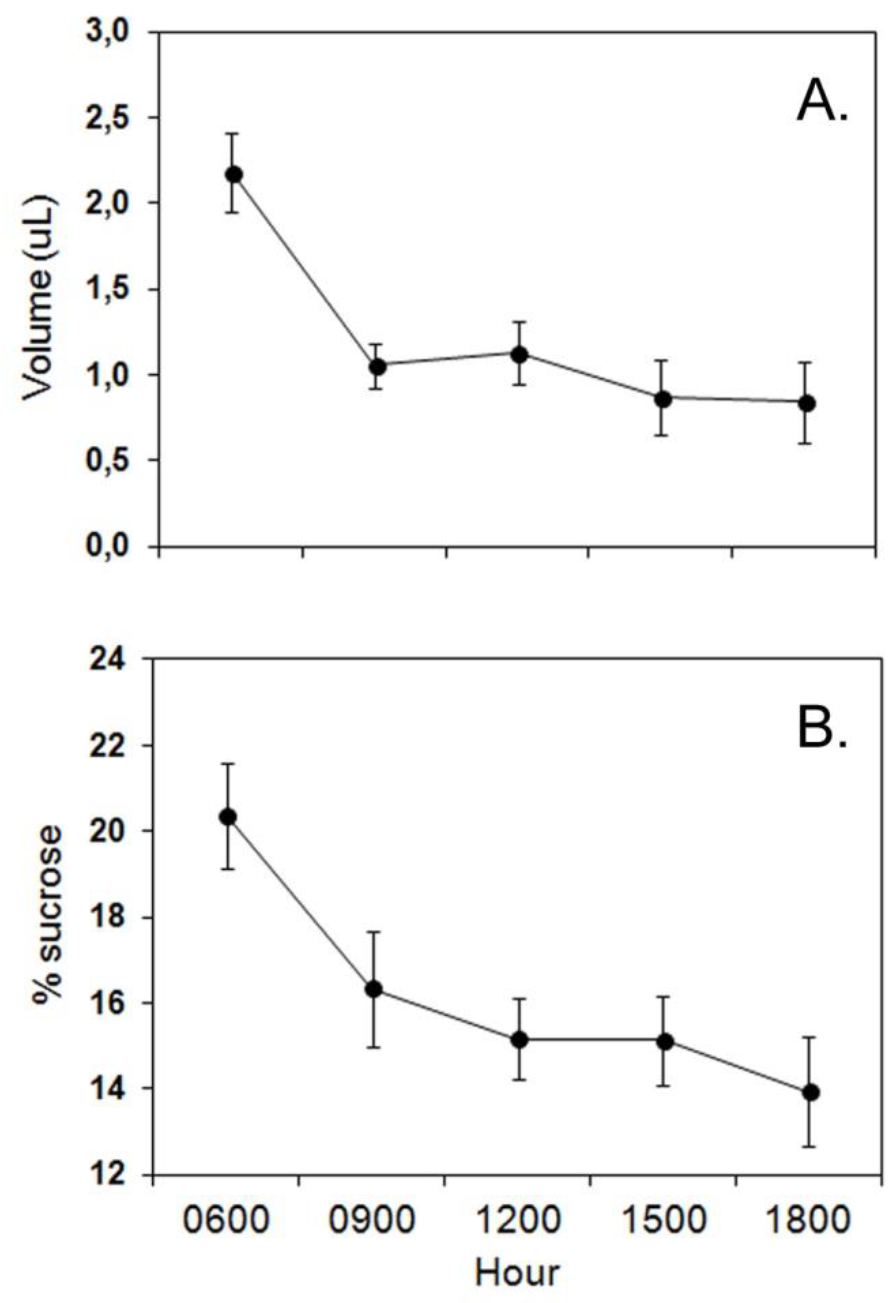
Diurnal pattern of nectar production of unvisited flowers of Pitcairnia heterophylla (Bromeliaceae) at a cloud montane forest, Costa Rica. Data are mean values from measurements on 18 flowers at 3-h intervals (vertical bars = 1 SE).

### Pollinators and floral visitation

Short-billed hummingbirds (Trochilinae) were the main visitors to *P. heterophylla* flowers. *Lampornis calolaemus* (Salvin, 1865) accounted for most of the recorded visits (78% Fig. 1D), followed by *Eupherusa eximia* (DeLattre, 1843), and *Selasphorus scintilla* (Gould, 1851). Peak activity in visitation pattern occurred between 1000 and 1300 h, comprising 61% of the recorded visits (Fig. 3). Only 46 visits to 97 flowers were recorded over 918 hours of camera monitoring and, the daily and hourly visitation rate per genet was rather low (0.6 visits day^-1^, mean = 0.05 visits hour^−1^; range=0.0−0.19). A nectar robber *(Diglossa plumbea,* Emberizidae) and a floral herbivore, the red-tailed squirrel *(Sciurus granatensis,* Sciuridae), were also recorded on inflorescences of the studied species.

**Fig. 3.**
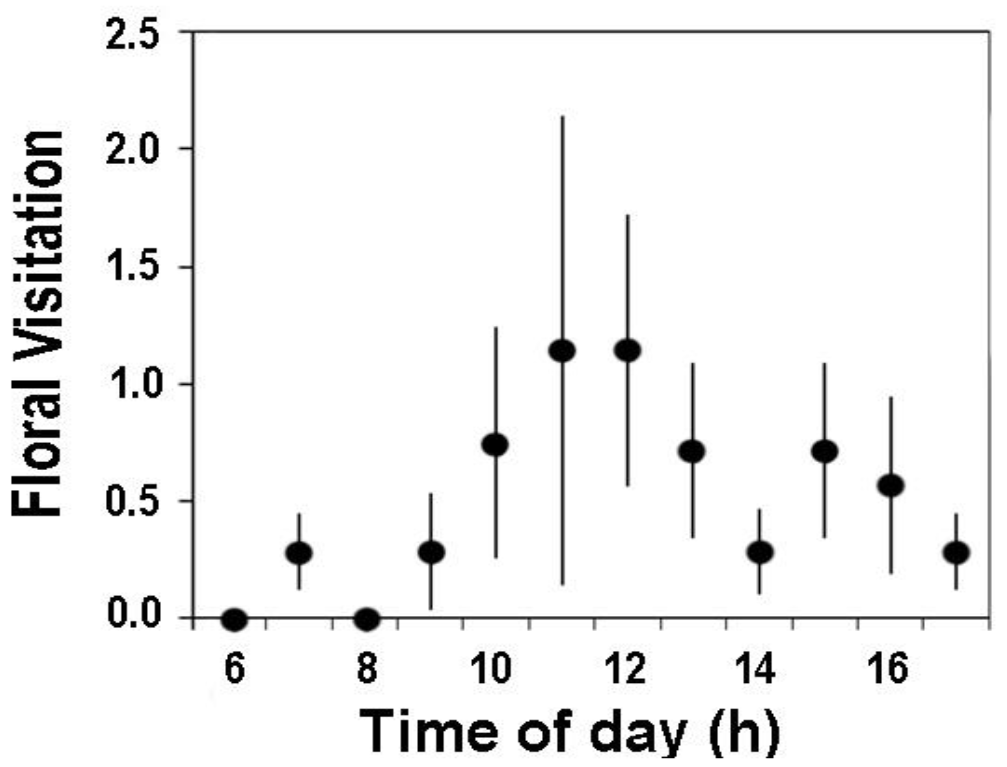
Rate of hummingbird visitation to flowers of Pitcairnia heterophylla (Bromeliaceae) at a cloud montane forest, Costa Rica. Data are average numbers of visits per hour (lines are ±1 SE) from the reproductive season of 2013 and comprising 918 observation hours on eight individuals. Total visits = 46.

## Discussion

Our results indicate that *P. heterophylla* is pollinated by hummingbirds and capable of high outcrossing rates but, at the same time, exhibits floral traits that promote selfing, which in turn suggests a mixed mating system. Mixed mating is a common phenomenon in biotic pollination systems (Goodwillie *et al.* 2005) and involves autonomous self-pollination and self-compatibility. A similar mixed mating system has been documented for other *Pitcairnia* species (Wendt *et al.* 2001, 2002, Fumero-Caban & Melendez-Ackerman 2007, Bush & Guilbeau 2009) and it seems a common reproductive strategy among epiphytic bromeliads (reviewed by Matallana *et al.* 2010).

Pollination by hummingbirds in *P. heterophylla* is congruent with the results from other *Pitcairnia* species, both epiphytic (Bush & Guilbeau 2009) and terrestrial (Wendt *et al.* 2001, 2002). The scentless flowers, with vivid scarlet and tubular corolla and nectar production in *P. heterophylla* fulfill the criteria of an ornithophilic plant (Willmer 2011), which was confirmed by the video recording of hummingbird visitors. However, *P. heterophylla* presents lower nectar production and sugar concentration, when compared to other hummingbird pollinated species (Krömer *et al.* 2008, Ramírez & Ornelas 2010, Fumero-Caban & Melendez-Ackerman 2012).

The low visitation rate recorded from *P. heterophylla* plants (0.6 visits day^−1^) at the study site contrast with reports from several hummingbird pollinated plants at a similar cloud forest ecosystem in Costa Rica (Feinsinger 1978, Bush & Guilbeau 2009). Local pollinator abundance and competitive interactions among hummingbird-pollinated plants in the community may influence the visitation rate and affect the reproductive success (Stiles 1975). In our case, *P. heterophylla* is the only representative of the genus in the studied site, thus it is unlikely that selfing might serve to reduce interspecific pollination as proposed by Matallana *et al.* (2010). Several sympatric hummingbird-pollinated species overlap their flowering period with that of *P. heterophylla* at the study site (A. Cascante-Marín, unpubl. data), thus selfing might counteract the potential deposition of pollen from ornithophilic species belonging to unrelated taxonomic groups. In such situation, the ability of *P. heterophylla* to deposit self-pollen on the stigma would prevent the contamination with foreign pollen.

Pollination success in *P. heterophylla* might also be affected by phenological events related to leaf shedding and flower anthesis at the individual level (Fig. 1B). The short inflorescence of *P. heterophylla* could potentially be concealed by the long linear photosynthetic leaves while in most other *Pitcairnia* species the longer (>20 cm) inflorescences lie above the leaves (Smith & Downs 1974). However, at the time of flowering, ramets of *P. heterophylla* have shed the linear leaves and exposed the inflorescences which become available to pollinators. Hummingbirds visually search for their food source; thus flowers must be exposed in order to attract them (Faegri & van der Pijl 1979). Reproductively, leaf shedding in *P. heterophylla* might function to increase flower visibility and pollinator visitation, as it has been suggested for some tropical dry forest trees which exhibit a similar phenological behavior (Janzen 1967).

Floral mechanisms that facilitate autonomous selfing are common among plants with specialized pollination systems (Fenster & Marten-Rodriguez 2007) and, in general, the evolution of selfing in plants has been interpreted as a means to attain reproductive assurance whenever pollination conditions are limiting (e.g., low pollinator or mate abundance) (Jain 1976; Holsinger 2000; Charlesworth 2006). Our results from the breeding system analysis of *P. heterophylla* indicated that the species has a relatively high potential to sire progeny by autonomous means (AFIf=0.79), a condition facilitated by self-compatibility, adichogamy and absence of herkogamy. It is likely that flowers of *P. heterophylla* self-pollinate as a way to cope with low hummingbird visitation and ensure their seed set. On the other hand, the lower seed production recorded in manually selfed versus manually crossed flowers constitutes evidence of reduced female fitness through seed discounting (Lloyd 1992). The potential effects of endogamy on plant fitness through tests on seed germination and plant growth between outcrossed and selfed progeny are needed to assert this supposed advantage of selfing in *P. heterophylla.*

Our findings with *P. heterophylla* are in congruence with the hypothesis of reproductive assurance when pollinator service is unpredictable (Levin 1972). Although we weren’t able to expose emasculated flowers to natural pollination conditions, our results fit such hypothesis as the addition of exogenous pollen significantly increased seed set. Also, in a similar scenario to the one we described above, where low pollinator visitation is common in hummingbird-pollinated and epiphytic bromeliads, outcrossing might be restricted and selfing may serve to increase reproductive success. Molecular analysis would probably reveal a highly inbred progeny in *P. heterophylla* as demonstrated in other epiphytic bromeliads with similar pollination systems (Cascante-Marín *et al.* 2006). The evolutionary advantage of selfing as a mechanism of reproductive assurance to compensate for pollinator limitation in the studied species needs to be corroborated through experiments that evaluate the effects on pollen and seed discounting *sensu* (Lloyd 1992) and fitness reduction of the progeny.

## Acknowledgments

The Guide and Scout Association of Costa Rica granted permission to conduct the research at the Iztarú Camp at Cerros La Carpintera. Gilbert Barrantes and Diego Ocampo (UCR) identified the hummingbirds. Eric J. Fuchs provided valuable comments to a previous manuscript. This work was supported by Vicerrectoria de Investigation at Universidad de Costa Rica (project 111-B2-041 awarded to ACM). The Agencia Espanola de Cooperation Internacional y de Desarrollo (AECI) provided funds to acquire video camera equipment (funds D/027406/09 and D/033858 to Universidad de Las Palmas de Gran Canaria and Universidad de Costa Rica).

